# Convergent Usage of Amino Acids in Human Cancers as a Reversed Process of Tissue Development

**DOI:** 10.1101/2021.03.18.436083

**Authors:** Yikai Luo, Han Liang

**Affiliations:** Graduate Program in Quantitative and Computational Biosciences, Baylor College of Medicine, Houston TX 77030, USA; Department of Bioinformatics and Computational Biology, The University of Texas MD Anderson Cancer Center, Houston, TX 77030, USA; Department of Systems Biology, The University of Texas MD Anderson Cancer Center, Houston TX 77030, USA

**Keywords:** Amino acid usage, Tissue development, Biosynthetic energy, Diagnostic biomarker

## Abstract

Genome and transcriptome-wide amino acid usage preference across different species is a well-studied phenomenon in molecular evolution, but its characteristics and implication in cancer evolution and therapy remain largely unexplored. Here, we analyzed large-scale transcriptome/proteome profiles such as TCGA, GTEx, and CPTAC and found that compared to normal tissues, different cancer types showed a convergent pattern towards using biosynthetically low-cost amino acids. Such a pattern can be accurately captured by a single index based on the average biosynthetic energy cost of amino acids, termed Energy Cost Per Amino Acid (ECPA). With this index, we further compared the trends of amino acid usage and the contributing genes in cancer and tissue development and revealed their reversed patterns. Finally, focusing on the liver, a tissue with a dramatic increase in ECPA during development, we found that EPCA represented a powerful biomarker that could distinguish liver tumors from normal liver samples consistently across 11 independent patient cohorts (AUROC = ~0.99) and outperformed any index based on single genes. Our study reveals an important principle underlying cancer evolution and suggests the global amino acid usage as a system-level biomarker for cancer diagnosis.

## Introduction

Amino acids are the basic building blocks of a cell. Coding sequences and gene expression profiles are two key factors determining the overall amino acid usage of a cell. Through analysis of the genomes or transcriptomes of many species, preferred amino acid usage is a well-studied topic in macroevolution. The universal trend of “Cost-Usage anticorrelation” suggests that the relative abundance of amino acids, quantified as the number of codons encoding a specific amino acid in the genome of a species, is mainly driven by their biosynthetic energy costs [1–5]. However, it remains unclear how amino acid usage of cancer cells deviates from normal tissues and evolve in different tumor contexts.

From an evolutionary point of view, cancer cells are characterized by a low degree of divergence from its tissue of origin, measured by the limited amount of somatic changes, which is in contrast to the macroevolution that happens across different taxa or even the microevolution existing between within-species individuals [6]. However, such trifling transformation does yield a wide range of phenotypic commonalities shared by distinct cancer types, including activated proliferative signaling, resistance to programmed cell death, induction of angiogenesis, and metastatic capability [7]. Among many theories proposed to understand such convergence, one appealing concept is that cancer cells bear a set of genomic, transcriptomic, and epigenomic features that can be summed up as “stemness,” [8–11] which in the context of ontogeny, defines the level of reprogramming/dedifferentiation of adult tissue cells. The underlying mechanistic links between cancer evolution and tissue development have been hinted at by the observations of frequent mutations leading to reactivation of stem cell-related pathways in cancer [12,13]. However, little effort has been made to examine a potential association between these two seemingly non-overlapping processes with respect to amino acid usage.

Characterizing the amino acid usage of cancer cells not only helps us understand the evolutionary constraints in the tumor microenvironment but may also have clinical utility. In recent years, tremendous efforts have been made to identify gene expression-based biomarkers for cancer diagnosis, outcome prediction, and treatment selection, but successful cases with proven clinical values are still limited [14–16]. One factor that determines the feasibility of such biomarkers in clinical practice, the robustness, is rarely satisfied, meaning that a threshold chosen based on limited data is usually not generalizable to unseen scenarios. In contrast to conventional biomarkers based on individual genes, the amino acid usage represents a holistic property of a cellular state. Therefore, there is a possibility that its related indices represent more robust biomarkers for clinical applications. To fill these knowledge gaps, here we performed a systematic analysis of the amino acid usage profiles across many cohorts of tumor and normal tissue samples.

## Results

### A convergence of amino acid usage across cancer types

Since gene expression levels are largely associated with amino acid usage in a cell, we first examined the gene expression patterns of 30 tissue types in the Genotype-Tissue Expression (GTEx) cohort [17] (Figure S1A) and 31 cancer types in The Cancer Genome Atlas (TCGA) cohort [18] (Figure S1B). Using the t-distributed stochastic neighborhood embedding (t-SNE)[19] projection, we found that samples of a common tissue origin largely formed a single cluster regardless of being normal or cancerous. In addition, cancer types with the same tissue origin, such as brain cancers (glioblastoma multiforme [GBM] and lower grade glioma [LGG]), kidney cancers (kidney renal clear cell carcinoma [KIRC] and kidney renal papillary cell carcinoma [KIRP]), lung cancers (lung adenocarcinoma [LUAD] and lung squamous cell carcinoma [LUSC]), and liver cancers (hepatocellular carcinoma [LIHC] and cholangiocarcinoma [CHOL]), tended to be mingled or closer to each other than to other cancer types. We observed similar patterns in two other large, pan-cancer cohorts, PCAWG [20], and MET500 [21] (Figure S1C and D). Consistent with previous studies [18,22], these results indicate that cancer cells largely retain their tissue-specific gene expression profiles.

To study whether this tissue-specific pattern holds for amino acid usage, we calculated the similarity of transcriptome-based amino acid usage by integrating the gene expression profiles and the amino acid frequencies of protein-coding genes (**Figure 1A**) and visualized their patterns in the same way. Similar to the strong tissue specificity observed in the gene expression analysis, we found that normal tissues of the GTEx cohort still had distinct amino acid usage patterns (Figure 1B). We further confirmed this result by co-clustering amino acid usage profiles of the Human Protein Atlas (HPA) cohort [23] with corresponding GTEx tissue types (Figure S2A). More intriguingly, samples of a multi-species multi-tissue cohort [24] were principally separated by tissue type rather than by species, suggesting that tissue-specific amino acid usage is highly conserved across mammals (Figure 1C).

**Figure 1.**
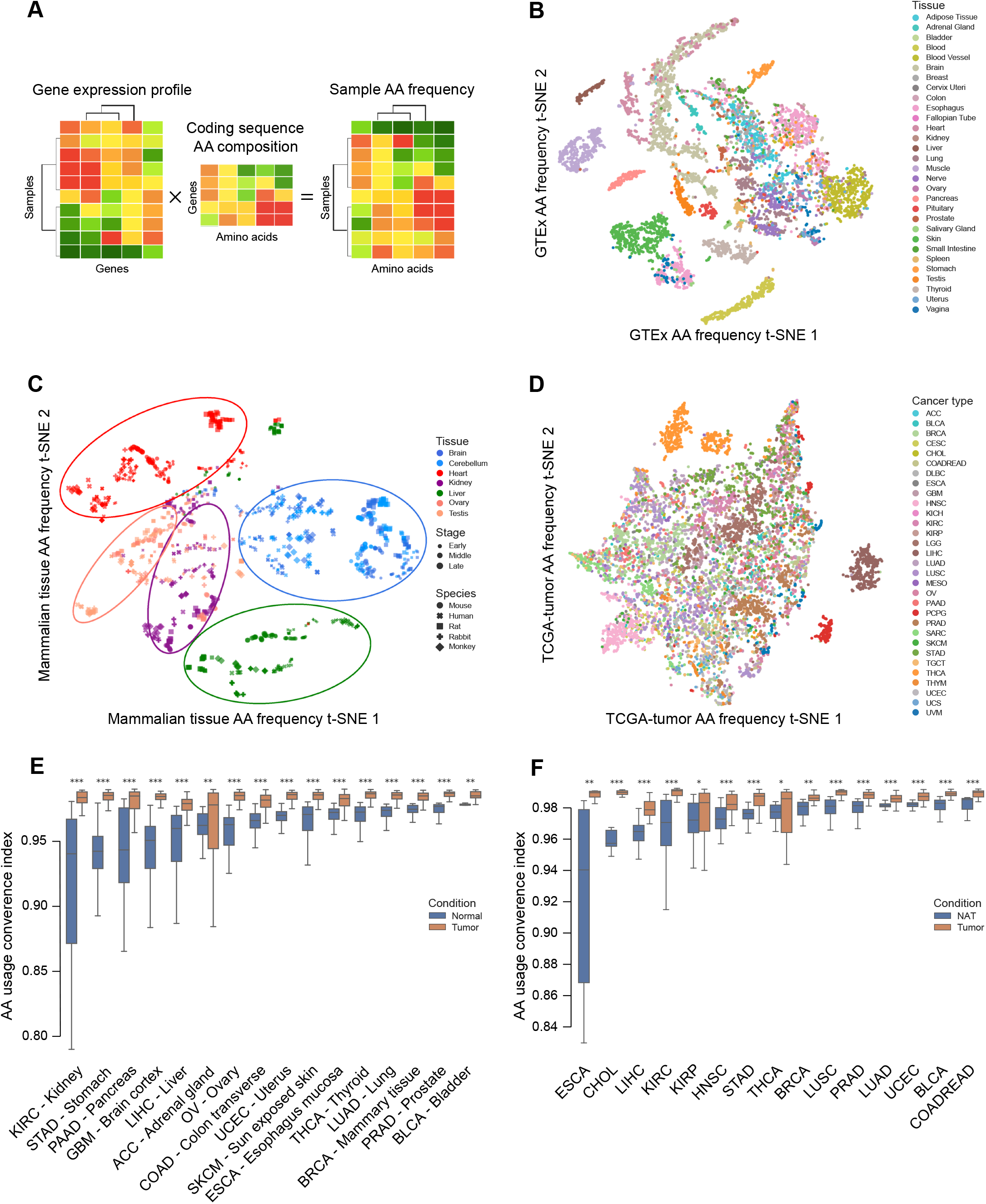
Pan-cancer convergence of transcriptome-based amino acid usage. **A**. Schematic diagram showing the computation of amino acid usage frequency based on the gene expression profile derived from an RNA-seq sample. t-SNE projection of the GTEx (**B**), developing mammalian tissue (**C**), and TCGA tumor samples (**D**) based on their amino acid frequency profiles. Samples are color-coded based on tissue or cancer types. Marker shapes correspond to species. Developmental stages were classified into three categories and indicated by marker size. All t-SNE projections were generated using sklearn TSNE, with perplexity as 30, learning rate as 200, and the number of iterations as 1,000. Comparison of amino acid usage convergence index between tumor samples and either matched down-sampled normal samples (**E**) or adjacent normal samples (**F**) across multiple cancer types. Box plots show the quartiles, and the whiskers indicate quartile ± 1.5 × interquartile range. A two-sided Mann-Whitney U-test was used to calculate the p-value. *p < 0.05, **p < 0.01, ***p < 0.001.

In sharp contrast to normal tissues, when clustered by amino acid usage, samples of different cancer types were much less separated and did not segregate on the basis of tissue origins (Figure 1D). To further confirm this observation, we clustered amino acid usage profiles of two other cancer cohorts, PCAWG and MET500, and observed a dramatic loss of tissue-specificity relative to the patterns observed in the gene expression-based analysis (Figure S1C and D and Figure S2B and C). To ensure that the detected pattern was not due to a disparity in sample size or unmatched tissue types, we leveraged a conservative GTEx-TCGA mapping to only include normal and tumor samples whose tissue origins are matched without ambiguity, then performed down-sampling within individual tissue-specific cohorts, and finally, applied t-SNE to redo a supervised clustering. The results remained the same for the comparison between down-sampled GTEx and TCGA samples (Figure S2D and E) as well as for that between TCGA tumor samples and the normal adjacent to tumor (NAT) (Figure S2F and G). This observation is important since, evaluating tumor purity and gene signatures, recent studies have shown that NAT samples reside in an intermediate state between healthy and tumor [25,26].

The observation that amino acid usage for cancer cells failed to preserve their distinct tissue origins raised two possibilities: (i) cancer cells evolved to possess highly stochastic amino acid usage profiles both within and between cancer types; or (ii) they went through convergence of amino acid usage, thereby losing the constraint of the original tissue specificity. To identify the correct hypothesis, we simply asked whether, in the 20-dimensional space (each dimension representing the frequency of specific amino acid), the distances between samples of different cancer types were shorter than those among samples of different normal tissues. Based on Pearson’s distance, for each sample, we defined an amino acid usage convergence index that measured its distance to all other samples of different tissue or cancer types. Through a comparative analysis of GTEx normal vs. TCGA tumor and TCGA NAT vs. tumor, we found that tumor samples showed significantly increased convergence than normal samples, a pattern consistently observed across all surveyed cancer types (Figure 1E and F). Furthermore, we compared the variations of amino acid frequencies across NAT samples and tumor samples of different cancer types based on the same set of standard deviations. Indeed, the extent to which amino acids are differentially used in tumors was markedly reduced than that in NATs (Figure S4A and B). Collectively, these results indicated a strong convergence rather than a stochastic transformation of amino acid usage across cancer types, supporting our second hypothesis.

### Cancer cells tend to use biosynthetically low-cost amino acids

To understand how such a convergent pattern occurs, we quantified the differential usage of each amino acid in tumors vs. normal tissues and found no highly consistent trend across cancer types in terms of increased or decreased usage (Figure S4C). However, when taking a higher view of the heatmap, structurally complex amino acids, such as tryptophan and cysteine, tended to be significantly depleted in most cancer types, whereas those with relatively simpler structures tended to be significantly enriched in a majority of cancers. Because the structural complexity of the amino acids correlates well with the energy cost of their biosynthesis [1], we hypothesized an association between the biosynthetic energy cost of amino acid and its usage tendency in cancers. Indeed, we observed a strong negative correlation between the biosynthetic energy cost and the net number of cancer types in which the usage of an amino acid was significantly increased (**Figure 2A**, Rs = −0.56, p = 0.01), suggesting that cancer cells prefer amino acids with a lower biosynthetic energy cost. We previously introduced two indices, ECP_Agene_, and ECPA_cell_, which quantify the average biosynthetic energy cost per amino acid for a gene and a cell (or a sample), respectively [27] (Figure 2B). ECP_Agene_ is based on the amino acid frequency encoded in a gene, and ECPA_cell_ considers the expression levels and amino acid frequencies of all the genes in a cell. A high ECPA value indicates that the gene or the cell tends to use biosynthetically expensive amino acids. We found that compared to NAT samples, ECPA_cell_ of the tumor samples became significantly lower for 9 out of the 15 tested cancer types, while no significantly opposite patterns were observed (Figure 2C). To confirm this pattern at the proteomic level, we extended these analyses to six cancer proteomics datasets from the Clinical Proteomic Tumor Analysis Consortium (CPTAC) [28] and others [29,30], covering five cancer types. Strikingly, in all the cases, proteins that were significantly up-regulated in tumor samples (log_2_FC > 0, FDR < 0.05) had significantly lower ECP_Agene_ than the proteins that were significantly down-regulated (log_2_FC < 0, FDR < 0.05) (Figure 2D). These results indicate that cancer cells reshaped their gene/protein expression programs to use biosynthetically inexpensive (or structurally simpler) amino acids, thereby losing their original tissue-specific amino acid usage profiles. Finally, we sought to test if our ECPA index is insensitive to the expression of genes with extremely high abundance, including those encoding certain housekeeping proteins as well as tissue-specific proteins. After removal of all genes that either encode cytoplasmic and mitochondrial ribosome proteins or rank within top 200 in median TPM of the same cancer type, we recalculated the ECPA index for each sample and found that the pattern of consistent decrease of ECPA_cell_ in tumor samples across multiple cancer types was almost perfectly reproduced (Figure S3).

**Figure 2.**
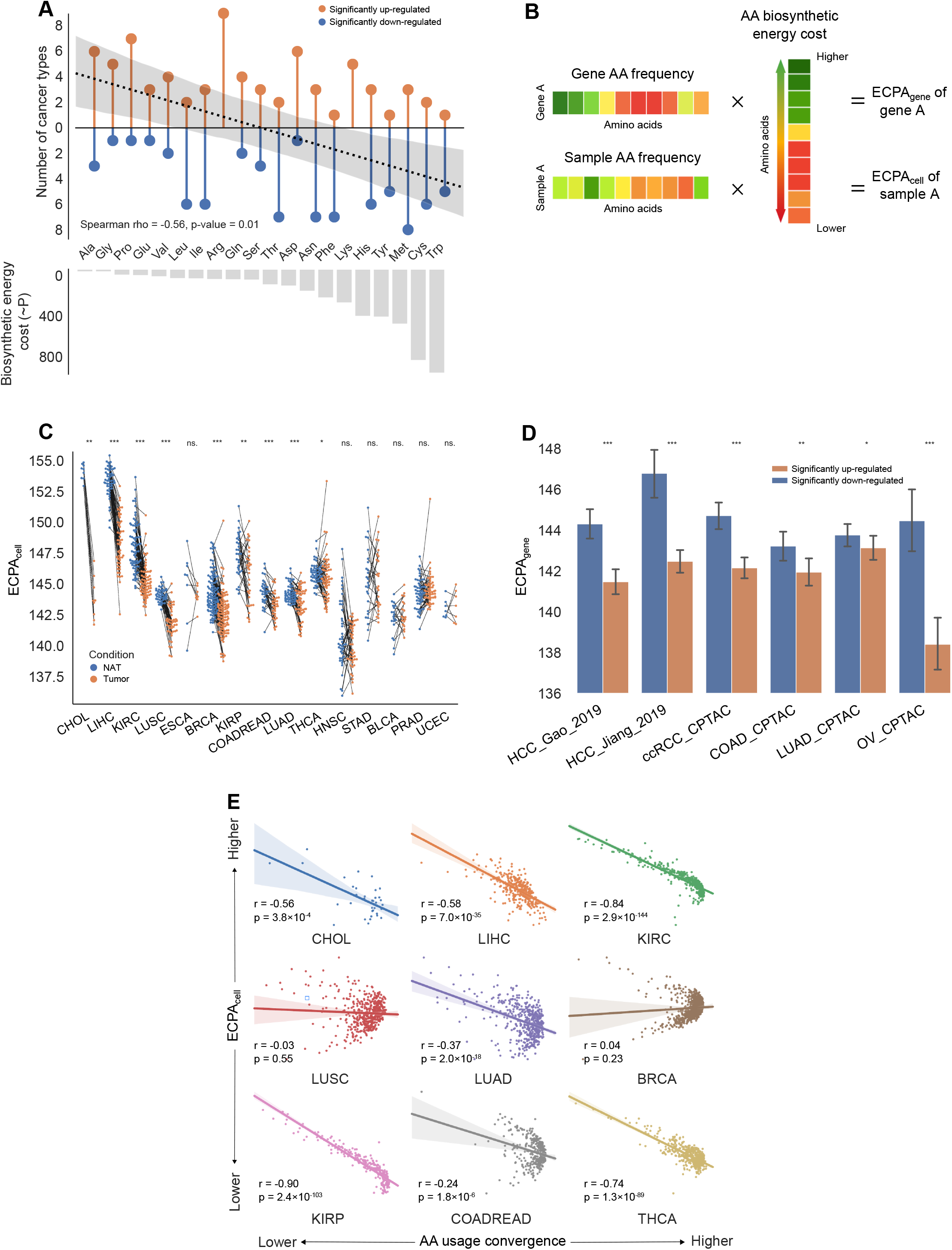
Amino acid usage preference in tumor evolution as quantified by ECPA_cell_. **A**. Correlation between the biosynthetic energy cost of an amino acid and the net number of cancer types with significantly increased usage across 20 amino acids. The net number is defined as the number of cancer types with significantly increased usage of the amino acid minus the number with significantly decreased usage. The colored region around the regression lines indicates a 95% confidence interval. **B**. Schematic diagram showing the computation of ECP_Agene_ and ECPA_cell_ based on the gene expression profile derived from RNA-seq data. **C**. ECPA_cell_ of tumor samples and matched normal tissue samples across TCGA cancer types. A paired two-sided Wilcoxon signed-rank test was used to calculate the p values. **D**. Bar plots showing ECP_Agene_ values of significantly down-and up-regulated proteins in several cancer proteomics datasets. Error bars denote 95% confidence intervals. A two-sided Mann–Whitney U-test was used to calculate the p values. **E**. Correlation between ECPA_cell_ and amino acid usage convergence index across samples in nine cancer types. The colored regions around the regression lines indicate 95% confidence intervals. *p < 0.05, **p < 0.01, ***p < 0.001.

We next tested whether the amino acid usage convergence level of a tumor was correlated with its ECPA_cell_. Indeed, we found a strong inverse relationship for seven out of the nine cancer types where ECPA_cell_ was significantly lower in tumors (Figure 2E). Thus, the more a tumor follows a convergent path to a common state of amino acid usage, the higher the bias it has toward using biosynthetically low-cost amino acids. These results also suggest that ECPA_cell_ is a simple, informative, interpretable index that effectively captures the overall preference of amino acid usage for a specific sample. Therefore, we focused on this index in subsequent analyses.

### Biosynthetically expensive amino acids are increasingly used during tissue development

To elucidate the underlying cause for the convergence of amino acid usage in cancer, we first sought to understand how tissue-specific amino acid usage patterns are established during development. Using the ECPA_cell_ index, we quantified the overall amino acid usage of liver and kidney tissues across different development stages in mammals, including humans, mice, rats, rabbits, and opossums. Intriguingly, both tissues showed an increasing trend of ECPA_cell_ along their developmental trajectories in all five mammals (**Figure 3A** and B). A closer inspection of the ECPA_cell_ trend lines led to two observations: i) key turning points of ECPA_cell_ in different species tend to happen at corresponding developmental stages; and ii) the rise of ECPA_cell_ in the liver takes concave trajectories while that in the kidney takes convex trajectories, suggesting that the establishment of high ECPA_cell_ status is driven by evolutionarily conserved synchronous molecular events that possess strong tissue specificity. To confirm this pattern, we collected another three independent RNA-seq datasets on mouse liver development and found a consistent ECPA_cell_ increase along the developmental paths in all three cases (Figure 3C-E).

**Figure 3.**
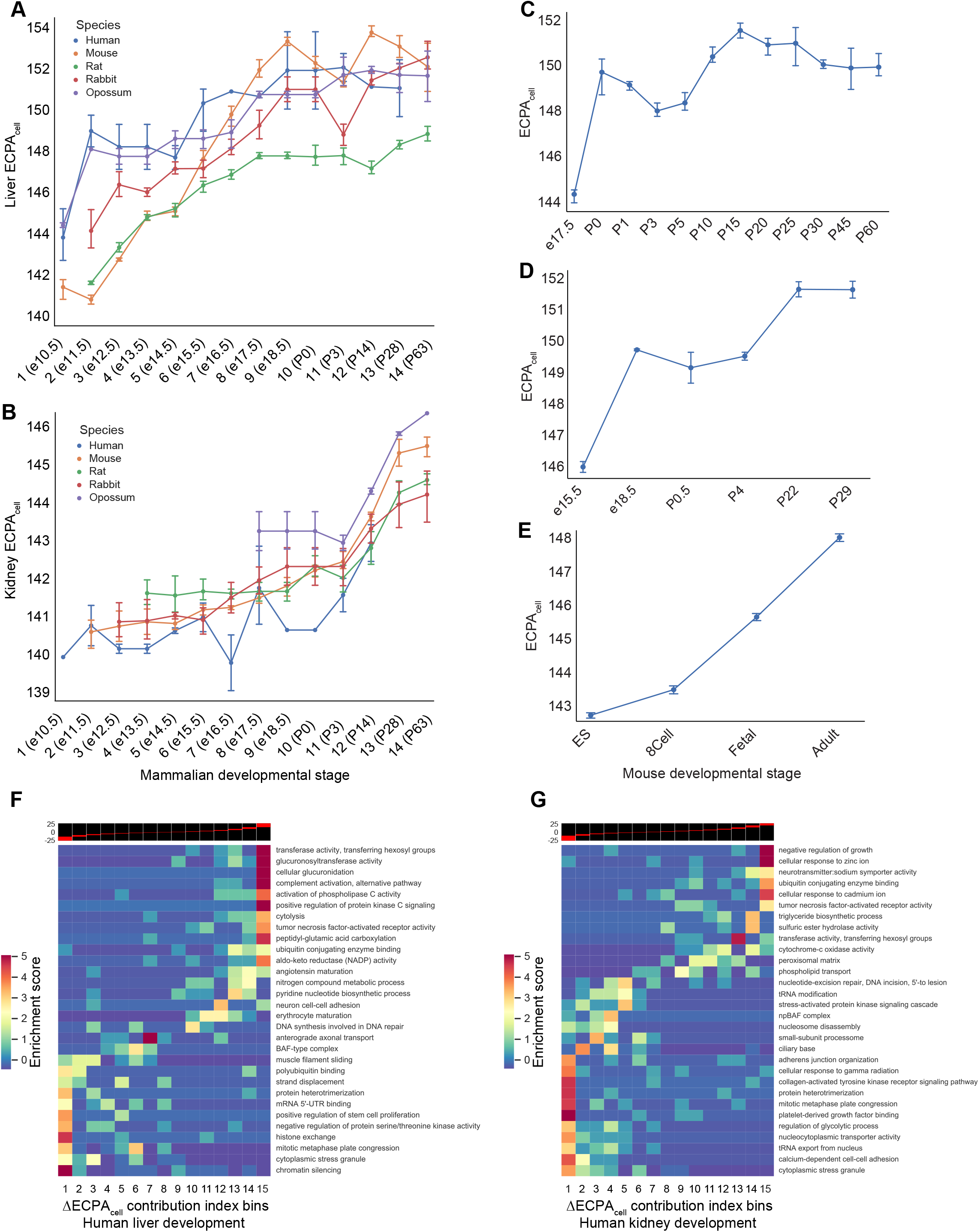
The increasing trend of ECPA_cell_ throughout mammalian organogenesis. Trend lines of ECPA_cell_ during the development of the liver (**A**), and the kidney (**B**) across five mammalian species. Developmental stages of non-mouse species correspond to the mouse stages shown in brackets. Error bars denote 95% confidence intervals. The trend line of ECPA_cell_ along the developmental trajectory of the mouse liver across three independent datasets (**C-E**). Error bars denote 95% confidence intervals. Heatmaps showing enrichment patterns of gene modules that contribute to ΔECPA_cell_ during the development of the human liver (**F**) and the human kidney (**G**). The red stripes embedded in the black background on top of each heatmap designate the range of ΔECPA_cell_ contribution index within every bin.

To pinpoint which gene modules are responsible for the tissue-specific build-up of a high ECPA_cell_ status, we first defined a “ΔECPA_cell_ contribution index” for each gene, which quantified the contribution of the gene to the global shift of ECPA_cell_ (see Materials and methods). We then divided all genes into 15 equal bins based on their index values and employed a mutual information-based enrichment identification algorithm called iPAGE [31] to detect the enrichment of these gene groups with well-established functional gene modules. We noted that genes contributing to the ECPA_cell_ increase were conserved among mammals but were tissue-specific. For the liver, the enriched modules included glucuronosyltransferase activity and complement activation (Figure 3F, Figure S5A, C and E); and for the kidney, the enriched modules included sphingolipid biosynthetic process and zinc/calcium ion homeostasis (Figure 3G, Figure S5B, D, and F).

Development-related cellular states that are instituted in adulthood can be prone to significant transformation or even complete collapse during aging [32]. To further understand how tissue-specific amino acid usage patterns alter when the tissue undergoes senescence, we gathered independent transcriptome profiles of aging livers and kidneys in humans, mice, and rats, and characterized the ECPA_cell_ patterns. Both tissues showed a stable pattern of high ECPA_cell_ status with reasonable fluctuations (Figure S6A-C). We concluded that tissue-specific, preferred usage of biosynthetically expensive (or structurally complex) amino acids, characterized by a high-ECPA_cell_ status, was gradually formed during development and remained largely unchanged in aging.

### Amino acid usage convergence of tumor follows a reversed path of tissue development

The strong convergence of amino acid usage across different cancer types is reminiscent of the “reverse-evolution” concept for tumorigenesis. As demonstrated above, this idea is well illustrated by the observation that there is a consistent decline of ECPA_cell_ in tumors, whereas there is a gradual increase of ECPA_cell_ during tissue development. To test the hypothesis that cancer evolution and tissue development move in opposite directions with respect to amino acid usage, we assessed whether the genes that boosted ECPA_cell_ in tissue development were overlapped with those that reduced ECPA_cell_ in tumors of the corresponding tissue origin and vice versa. Following the same method of computing ΔECPA_cell_ contribution index for tissue development, we measured the contribution of individual genes to ΔECPA_cell_ in cancer evolution for three cancer types for which gene expression profiles of normal developing tissues are available, namely LIHC, KIRC, and KIRP. Based on their contributions to ΔECPA_cell_ in either development or tumorigenesis, we divided individual genes into four quadrants with zero as the cutoff. We then used Fisher’s exact test to analyze the overlap of developmental ΔECPA_cell_-positive-contributing genes with tumorigenic ΔECPA_cell_-negative-contributing genes and vice versa. We observed that genes indeed tended to make opposite contributions to ΔECPA_cell_ in tumorigenesis and tissue development (**Figure 4A-C**, Fisher’s exact test, LIHC, p = 1.6×10^−156^; KIRC, p = 1.9×10^−39^; KIRP, p = 8.9×10^−30^). Furthermore, for the genes reducing ECPA_cell_ in tumorigenesis and increasing ECPA_cell_ in development, their ΔECPA_cell_ contribution index in these two processes were significantly negatively correlated (Figure 4D-F).

**Figure 4.**
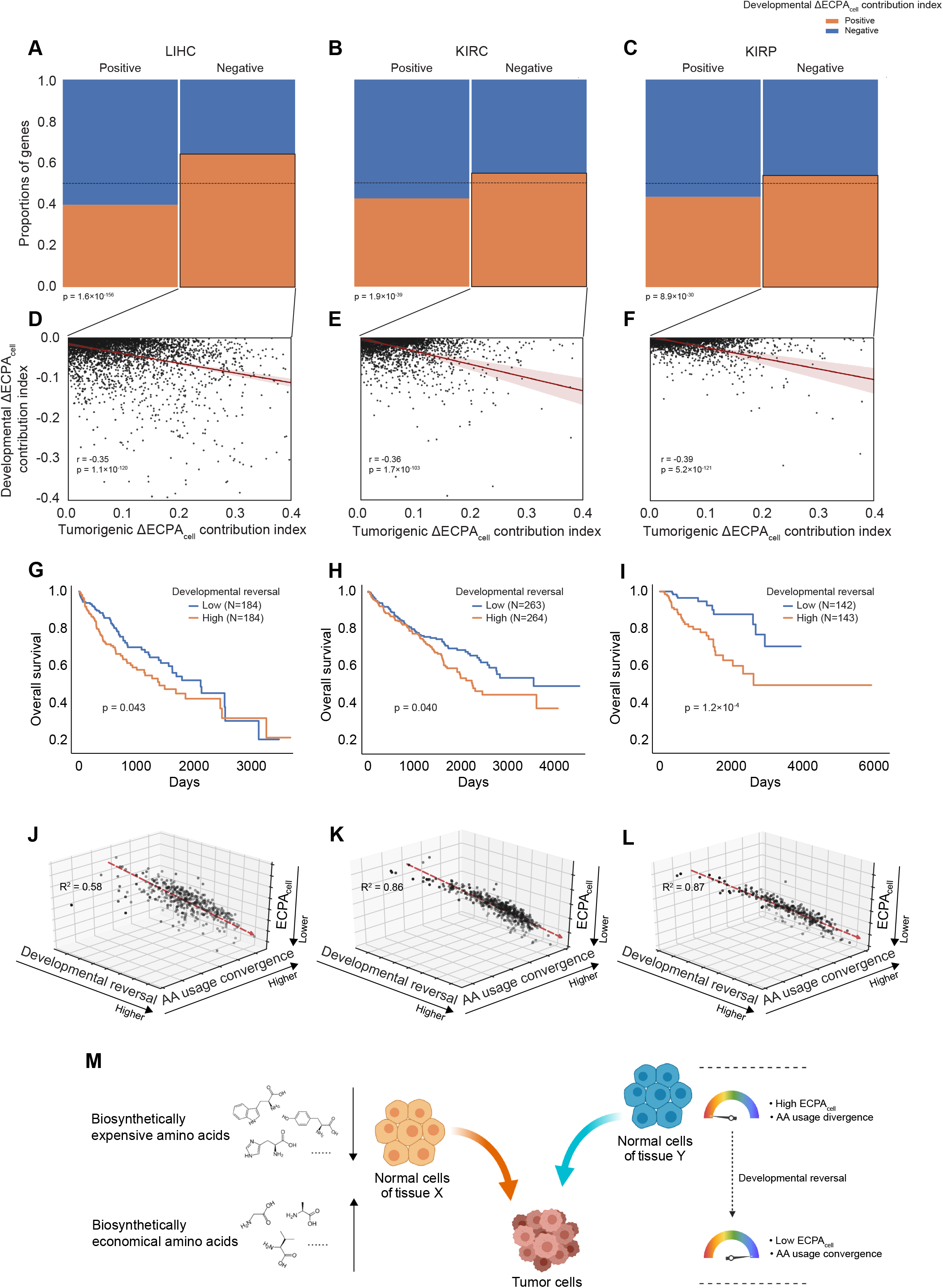
A proposed model unifying developmental reversal, amino acid usage convergence, and ECPA_cell_ decline of cancer samples. Stacked bar plots showing the proportion of genes that positively or negatively contribute to ΔECPA_cell_ in either tumorigenesis or development for LIHC-liver (**A**), KIRC-kidney (**B**), and KIRP-kidney (**C**). Scatter plots showing, for genes with negative ΔECPA_cell_ contribution index in tumorigenesis and positive ΔECPA_cell_ contribution index in tissue development, scaled ΔECPA_cell_ contribution index in tumorigenesis versus scaled ΔECPA_cell_ contribution index in tissue development for LIHC-liver (**D**), KIRC-kidney (**E**), and KIRP-kidney (**F**). Colored regions around the regression lines indicate 95% confidence intervals. Kaplan-Meier plots show the overall survival for patients with LIHC (**G**), KIRC (**H**), or KIRP (**I**) stratified by developmental reversal index into two equal groups, respectively. The p values were calculated from two-sided log-rank tests. Multivariate linear regression of ECPA_cell_ with developmental reversal index and amino acid usage convergence index as dependent variables for LIHC-liver (**J**), KIRC-kidney (**K**), and KIRP-kidney (**L**). **M**. Cartoon depicting a conceptual model in which cancer evolution is accompanied by the convergence of amino acid usage and decrease of ECPA_cell_, which is a reversal of the tissue development process.

While the gene-level analyses above were possibly hindered by the fact that cancer progression is highly heterogeneous even within the same cancer type [33,34], we can expect that a sample-level analysis would be more efficient to detect potential reverse relationships between cancer evolution and tissue development regarding amino acid usage. To this end, we defined the “developmental reversal index” for each tumor sample, which quantifies how strongly its gene expression pattern reversed what was instituted in tissue development. Specifically, we first calculated the gene-expression fold change of each tumor sample in terms of that averaged over the adjacent normal samples in order to measure the transcriptomic shift during tumorigenesis. We then measured the strength of anti-correlation between such a shift and the expression changes of the same gene set along the developmental trajectories of matched tissues (see Materials and methods). Interestingly, using this index to stratify cancer patients in terms of overall survival time, we found that a higher developmental reversal value was consistently associated with a worse prognosis (Figure 4G-I), suggesting that more aggressive tumors tend to have gene expression profiles more reversed in the tissue development trajectory.

Finally, we employed a multivariate linear regression model to clarify the associations between how biased a tumor sample tends to be in using biosynthetically inexpensive amino acids (represented by ECPA_cell_), how far it travels on the path of amino acid usage convergence relative to other cancer types (represented by amino acid usage convergence index), and how strongly its gene expression pattern reversed from what was instituted in tissue development (represented by the developmental reverse index). Remarkably, both the convergence level and the developmental reversal level were strongly anti-correlated with ECPA_cell_ across cancer types (Figure 4J-L). We, therefore, put forward an integrated model in which cancer cells initiated from distinct tissue origins converge into a common state favoring the use of biosynthetically inexpensive amino acids through reversed paths of tissue development (Figure 4M).

### The amino acid usage index, ECPA_cell_, is a robust biomarker for liver cancer diagnosis

Among different cancer types in our ECPA_cell_ analysis, the difference between liver normal and liver tumor samples was striking, making this tissue stand out from others (Figure 2C). Indeed, by quantifying the downward shift of ECPA_cell_ (ΔECPA_cell_) between tumor and the matched NAT pairs, the top two cancers were CHOL and LIHC, both of which originate from the liver (**Figure 5A**). We suspected that such a striking pattern could be attributed to liver-specific gene expression. To test this, we calculated ECPA_cell_ of both GTEx normal samples and TCGA NAT samples based only on tissue-specific genes [35] and ranked the tissues by their average ECPA_cell_. Indeed, the liver ECPA_cell_ level was higher than almost all other tissues (Figure 5B and C) (although the pancreas showed an even higher ECPA_cell_ according to the GTEx samples, the pattern did not hold for TCGA NAT samples). Of note, while the sample size of LIHC-NAT was as large as 50, the variation of their ECPA_cell_ based on tissue-specific genes was low. Furthermore, a comparison of the developmental ECPA_cell_ trend lines for different human tissues revealed that a fast and early build-up of a high-ECPA_cell_ status only existed for the liver (Figure 5D). We observed similar patterns in other mammals as well (Figure S7A-D). These results suggest that during development, the liver acquires a very high ECPA_cell_ state, and the liver-specific genes are the underlying contributing factor.

**Figure 5.**
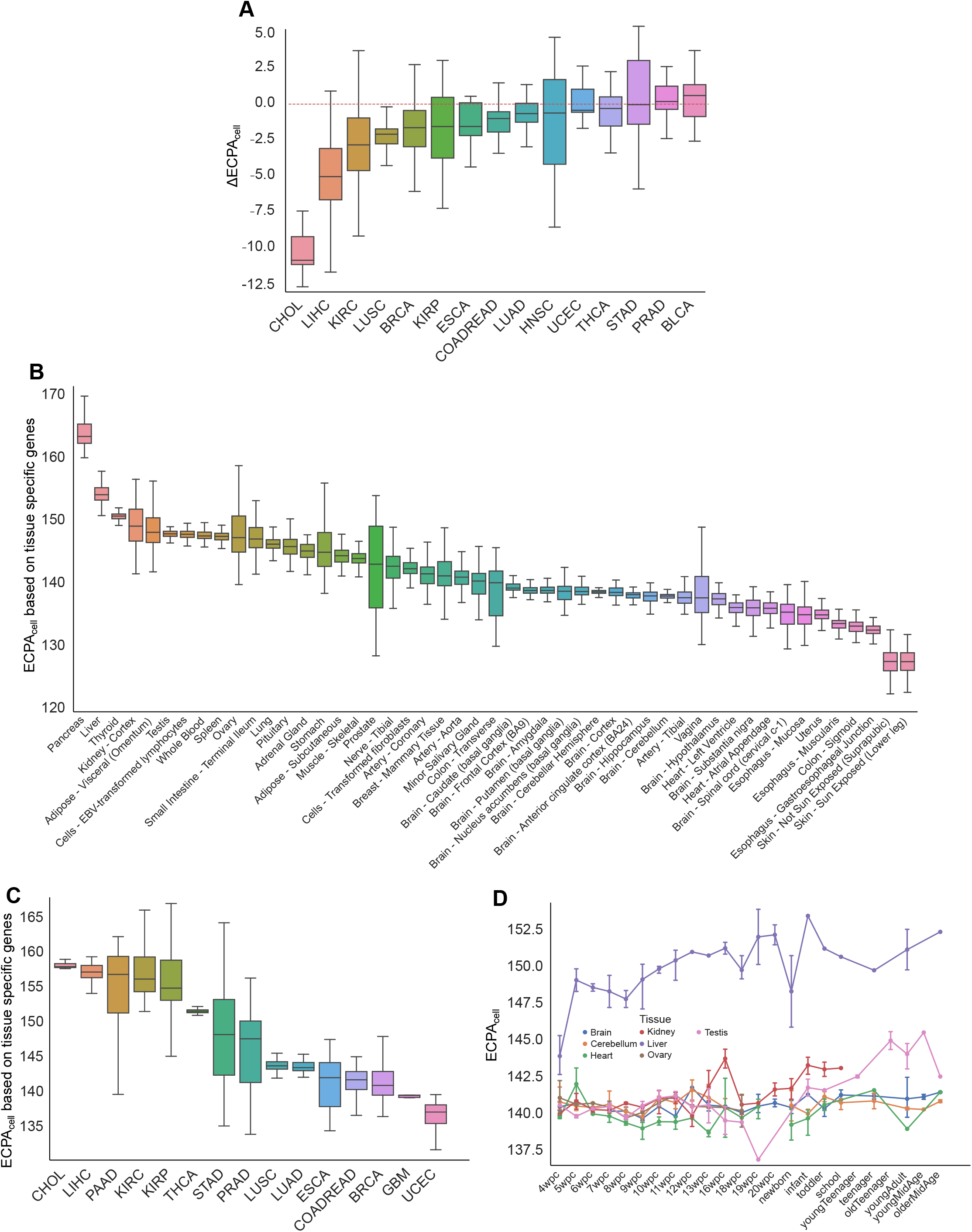
The liver shows the most dramatic ECPA_cell_ reduction in tumorigenesis. Distributions of ΔECPA_cell_ between tumor samples and paired NAT samples across multiple cancer types (**A**), tissue-specific genes-based ECPA_cell_ of normal samples across multiple tissues (**B**), tissue-specific genes-based ECPA_cell_ of adjacent normal samples across multiple cancer types (**C**), ranked by the median values. The box plots show the quartiles. The whiskers indicate quartile ± 1.5× interquartile range. The horizontal dashed line indicates the level of ΔECPA_cell_ = 0. **D**. Trend lines of ECPA_cell_ of multiple tissues across human developmental stages. Error bars denote 95% confidence interval. wpc, weeks post conception.

Given (i) the extremely high ECPA_cell_ level of liver tissue, and (ii) the dramatic difference between liver tumor and matched normal samples, we speculated whether ECPA_cell_ could be utilized as a novel biomarker for detecting liver cancer. To this end, we first collected 11 independent liver-cancer RNA-seq datasets (including TCGA LIHC and CHOL) where matched tumor and adjacent normal biopsies were simultaneously collected, thereby enabling a direct comparison of ECPA_cell_ between these conditions. In all cases, the tumor samples showed significantly reduced ECPA_cell_ with large effect sizes (**Figure 6A**).

**Figure 6.**
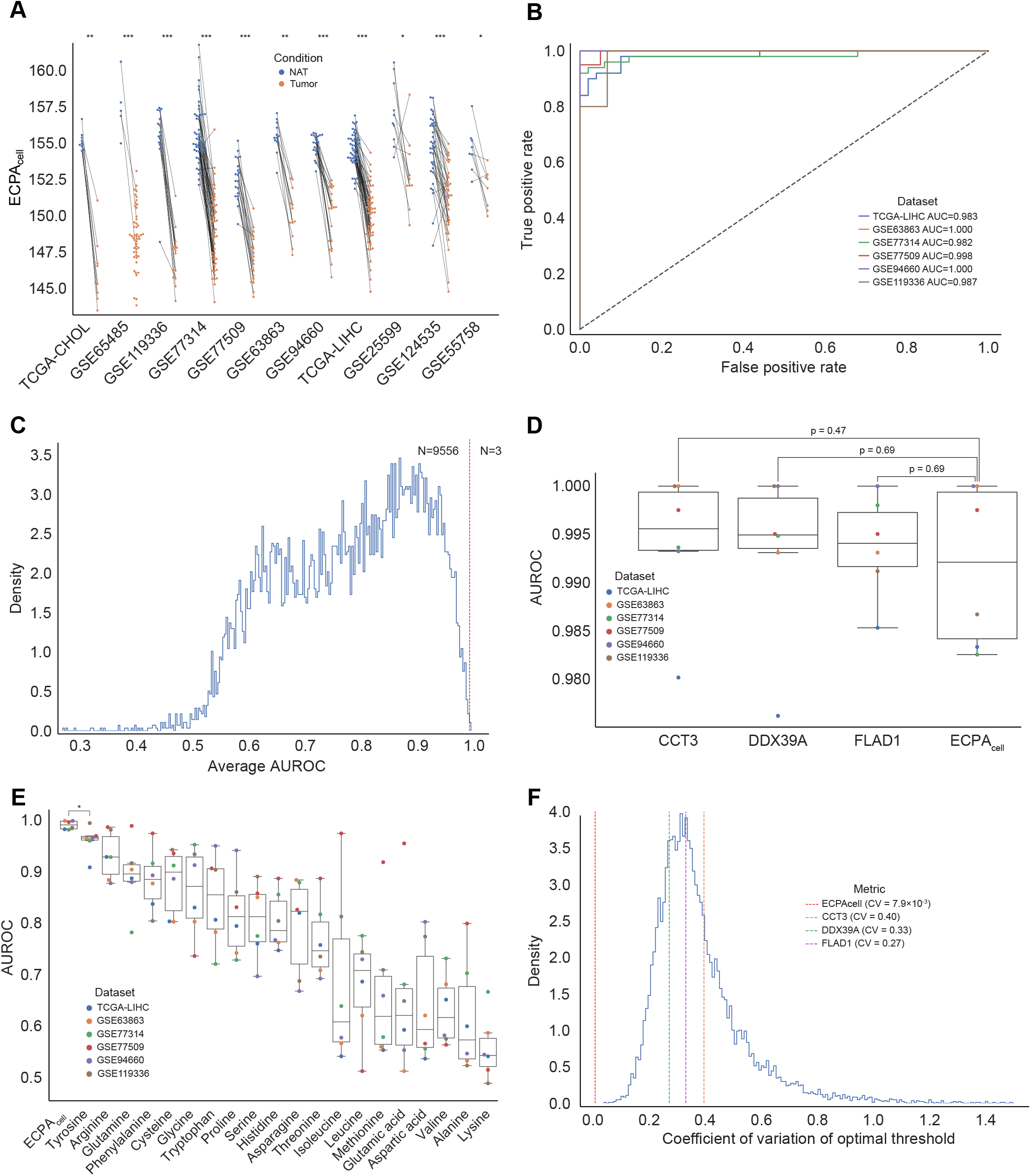
ECPA_cell_ is a robust diagnostic biomarker for liver cancer. **A**. ECPA_cell_ of tumor samples and matched normal tissue samples in 11 independent RNA-seq datasets of liver cancer and their matched normal samples. A paired two-sided Wilcoxon signed-rank test was used to calculate the p values. **B**. ROC curves of ECPA_cell_ as a diagnostic biomarker in six independent liver cancer cohorts with sample size ≥12. Dashed lines indicate the lines of identity. ROC, receiver operating characteristic; AUC, area under the ROC curve. **C**. Histogram showing the distribution of the average AUC across the six cohorts for tumor-normal segregation using the mRNA expression level of each of the 9,559 detectable genes. The vertical dashed line corresponds to the average AUC of ECPA_cell_. **D**. Box plots showing the AUC of the top four metrics, including three genes and ECPA_cell_, in discriminating tumor samples from normal samples across the six cohorts. A paired two-sided Wilcoxon signed-rank test was used to calculate the p values. **E**. Box plots showing the AUC of ECPA_cell_ and the frequency of each amino acid in detecting tumors across the six cohorts. The box plots show the quartiles. The whiskers indicate quartile ± 1.5× interquartile range. A paired two-sided Wilcoxon signed-rank test was used to calculate the p values. **F**. Histogram showing the distribution of coefficients of variation (CV) of the optimal thresholds in using individual genes for tumor-normal segregation. The vertical red dashed line indicates the CV of ECPA_cell_. Vertical lines in three other colors indicate the CV of three genes whose average AUCs are higher than ECPA_cell_. *p < 0.05, **p < 0.01, ***p < 0.001.

To evaluate more rigorously the capacity of ECPA_cell_ to serve as a diagnostic marker in discriminating liver tumors from normal tissues, we employed the area under the receiver operating characteristic curve (AUROC) as a performance metric. To ensure the robustness of our analyses, we only included six datasets with sample size ≥12. The ECPA_cell_ index was able to separate tumor vs. normal samples with very high ROC scores (median value = 0.993, range = 0.982 - 1.00, Figure 6B). To compare the predictive power of ECPA_cell_ relative to individual gene-based biomarkers, we calculated the average AUROC of all detectable genes across the six datasets and assessed their performance in the same way. Among 9,559 genes assessed, only three genes (*CCT3*, *DDX39A*, and *FLAD1*) showed slightly better performance than ECPA_cell_ (0.992), but none of them had statistically significant superiority (Figure 6C and D). In addition, ECPA_cell_ showed significantly higher discriminating power than the usage of any single amino acid (Figure 6E). Along with accuracy, a key feature of a successful biomarker is its robustness. To assess this feature, we computed the coefficient of variation (CV) for the optimal thresholds of ECPA_cell_ and individual genes across different datasets as an indicator of robustness. ECPA_cell_ showed exceptionally high robustness with its CV as low as 7.9×10-3, about 5× smaller than the lowest CV of any single gene-based biomarker (Figure 6F). Notably, the three genes that had a statistically insignificant advantage over ECPA_cell_ by AUROC had extremely unstable optimal cutoffs among different datasets, suggesting their limited power in detecting liver cancer across diverse clinical scenarios. Collectively, these results suggest that, as a system-level feature capturing the global usage of amino acids in a sample, ECPA_cell_ represents a promising biomarker for liver cancer diagnosis, and possesses both high accuracy and exceptional robustness.

## Discussion

Here we performed a systematic analysis on transcriptome and proteome-based amino acid usage across a broad range of cancer types. Using a previously introduced index, ECPA_cell_, our results revealed, for different tumors, a convergent pattern toward a cellular state of using more biosynthetically low-cost amino acids. In parallel, we studied the amino acid usage in the developmental trajectories of multiple organs and uncovered diverse paths into a tissue-specific high-ECPA_cell_ status that were evolutionarily conserved across mammals. Thus, a reverse relationship existed between cancer evolution and tissue development, which can be viewed as reminiscent of the widely accepted concept of the cancer cell “stemness.” Furthermore, given the long-standing parallels between phylogeny and ontogeny [36], supported by recent evidence [24,37,38], it would be reasonable to interpret cancer evolution as a reversed process of not only the development of an organism or its tissues but also the evolution of species. It has been argued that one key mechanism adopted by cancer cells to obtain fitness in spite of the diversity of the microenvironments is to unleash the force that is suppressed in multicellular organisms but is borne by unicellular organisms that are at the very bottom of the evolutionary hierarchy [39–43]. Thus, amino acid usage, a key aspect of cellular metabolism, may provide a unique perspective to understand the fundamental principles governing cancer progression, tissue development, and macroevolution, three evolutionary processes on different scales.

With the advances in transcriptome profiling technology, gene expression-based biomarkers have attracted wide attention for tumor detection and patient stratification. However, due to the high heterogeneity of cancer and intrinsically stochastic nature of gene expression, biomarkers based on either a single gene or a set of genes tend to suffer from numerical instability, thereby performing poorly. As demonstrated for liver cancer diagnosis, our ECPA_cell_ index represents a system-level biomarker that has at least three remarkable advantages. First, ECPA_cell_ captures a global cellular state by retaining the entire transcriptome as its information source, thereby conferring unparalleled robustness. Second, ECPA_cell_ was derived *de novo* from the gene expression profile of a sample, thus independent of external reference, which might introduce large noise predominantly attributable to batch effect. Third, in contrast to data-driven metrics, ECPA_cell_ has a well-defined biological meaning, the biosynthetic energy cost of amino acids. Because of these properties, ECPA_cell_ is an extremely robust diagnostic biomarker for liver cancer with a nearly constant threshold for tumor-normal segregation. Further efforts are warranted to assess the utility of this index in other cancer types and clinical applications.

## Materials and methods

### Data acquisition and processing

We obtained the gene-level expression values (e.g., fragments per kilobase per million [FPKM] or transcripts per million [TPM]) of the TCGA cancer sample cohorts, the GTEx normal tissue cohort, and the MET500 metastatic tumor cohort, from the Xena data portal (https://xenabrowser.net/datapages/); the HPA cohort from the HPA data portal (http://www.proteinatlas.org/); and the PCAWG cohort from the ICGC data portal (https://dcc.icgc.org/releases/PCAWG/transcriptome/). We also obtained the gene expression values of the mammalian tissue development cohorts from ArrayExpress (https://www.ebi.ac.uk/arrayexpress/) under the accession IDs E-MTAB-6769 (chicken), E-MTAB-6782 (rabbit), E-MTAB-6798 (mouse), E-MTAB-6811 (rat), E-MTAB-6813 (rhesus macaque), E-MTAB-6814 (human), and E-MTAB-6833 (opossum); and two independent RNA-seq datasets of mouse liver development from the Gene Expression Omnibus (GEO) under the accession IDs GSE58733 and GSE58827, as well as from ArrayExpress under the accession ID E-MTAB-2328. Finally, we obtained RNA-seq datasets of aging mouse liver and kidney from GEO under the accession ID GSE132040.

To convert gene-level FPKM values to TPM [44] values for a gene g_*i*_ in a sample s_*i*_, we used the formula:

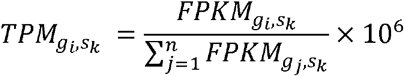

where the denominator on the right side is the sum of FPKM values of all the genes for an individual sample.

We downloaded raw RNA-seq fastq files of human liver cancer from GEO under the accession IDs GSE65485, GSE119336, GSE77314, GSE77509, GSE63863, GSE94660, GSE25599, GSE124535, and GSE55758; files of aging rat liver from the Sequence Read Archive (SRA) under the accession ID PRJNA516151, and files of TCGA LIHC and CHOL cohorts from the GDC Data Portal (https://portal.gdc.cancer.gov/). MultiQC [45] was used to assess the quality of the sequencing files and the performance of the preprocessing steps. Transcript-level abundances were quantified by Salmon [46] using the GRCh38 transcriptome as the reference. Gene-level TPM values were aggregated from transcript-level TPM values by tximport [47].

We obtained the proteomics datasets of KIRC, COAD, LUAD, and OV patient cohorts from the CPTAC data portal (https://cptac-data-portal.georgetown.edu/). We obtained two proteomics datasets of liver cancer from the NODE data portal (https://www.biosino.org/node/index/) and the CNHPP data portal (http://liver.cnhpp.ncpsb.org/), respectively.

### Calculation of transcriptome-based amino acid usage

We used the following formula to compute the amino acid frequency matrix given an RNA-seq dataset (see also Fig. 1a):

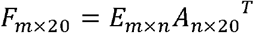

where *E* is a matrix of genes g_1_, g_2_, g_*n*_ by samples s_1_, s_2_, s_*m*_ with entries as TPM values, and A is a matrix of genes g_1_, g_2_, g_*n*_ by amino acids a_1_, a_2_, a_20_ with entries as relative frequencies of amino acids computed using the protein sequences annotated in the Swiss-Prot and TrEMBL databases hosted by the UniProt website (https://www.uniprot.org/). When a gene has multiple isoforms, we used its canonical sequence, as defined by UniProt based on criteria such as transcript length, relative abundance, and evolutionary conservation, in our analyses. We also repeated our analyses using transcript-level TPM data, where all isoforms annotated by ENSEMBL were included and had nearly identical results.

### Variation analysis of amino acid usage for TCGA samples

To illustrate the variation of amino acid usage of NAT samples from different tissues, we computed z-scores based on the average frequencies for individual amino acids across tissues. To compare these with the variations in amino acid usage of tumor samples across cancer types, instead of using *de novo* standard deviations to compute z-scores, we used the set of standard deviations derived for the NAT samples to obtain z-scores for the tumor samples. We used hierarchically clustered heatmaps with Euclidean distance as the distance metric to visualize the tissue-specificity of amino acid usage. To identify differential amino acid usage between tumor and NAT samples, we performed the Wilcoxon rank-sum test for frequencies of individual amino acids using paired tumor and NAT samples and used an FDR-adjusted p-value of 0.05 as the threshold for significance. Similarly, a hierarchically clustered heatmap was used to display amino acid de-regulation patterns across cancer types.

### Calculation of ECP_Agene_ and ECPA_cell_

We calculated two indices of amino acid usage, ECP_Agene_, and ECPA_cell_, representing the average biosynthetic energy cost per amino acid of a gene and a cell, respectively, as described previously [27]. Briefly, the biosynthetic costs of amino acids are based on the amount of high-energy phosphate bond equivalents required for amino acid biosynthesis in yeast and are normalized by amino acid decay rates (the biosynthetic costs of amino acids are highly correlated between different species). We then calculated ECP_Agene_ and ECPA_cell_ by multiplying the biosynthetic energy costs with the relative amino acid frequency of a gene or a cell (sample).

### Quantification of amino acid usage convergence for TCGA samples

To quantify the similarity of NAT or tumor samples in the TCGA cohort in terms of their amino acid usage patterns, we applied the Pearson’s distance metric to the amino acid frequency profiles, derived as described above. We also employed the Spearman rank correlation coefficient as an alternative metric and obtained the same results. Specifically, to capture the convergent pattern of amino acid usage across cancer types, we defined, for a sample s_*i*_ of cancer type *X*, the amino acid usage convergence index as:

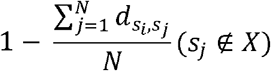

where 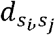 is the Pearson’s distance between sample s_*i*_ from cancer type *X* and sample s_*j*_ not from cancer type *X*.

### Calculation of ECPA_cell_ contribution index

To estimate the contribution of individual genes to the alteration of ECPA_cell_ in a specific biological process, we considered both how different the ECPA of a gene is from the baseline ECPA_cell_, as well as how much its expression level has changed. Formally, we defined the ΔECPA_cell_ contribution index of a gene *g*_*i*_ as:

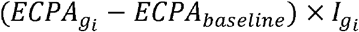

where 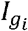 is an importance score that describes the extent of deregulation of *g*_*i*_. In tumorigenesis, we employed the log_2_ fold-change of average expression level between tumor and NAT samples as the importance score. In tissue development, we employed a different importance score that was not based on binary comparison as in tumorigenesis since the nature of the dataset is time-course measurements. Specifically, we applied an R package designed for transcriptomic time courses, maSigPro [48], to build a polynomial regression model (degree = 3) for each gene using its expression level as the response variable and the log-transformed post-conception days as the independent variable. Such models yielded the goodness-of-fit (R^2^) values that were then signed by the corresponding Spearman correlation coefficients and were finally used as the importance score.

### Pathway analysis of ECPA_cell_ contribution in mammalian tissue development

We employed an information-theoretic framework [31] to reveal gene modules or regulatory pathways that were enriched in genes with a significant contribution to the increase of ECPA_cell_ during tissue development. First, we focused on down-regulated genes with lower-than-baseline ECP_Agene_ and up-regulated genes with higher-than-baseline ECP_Agene_, both of which could contribute to the increase of developmental ECPA_cell_. Second, we distinguished these two groups of genes by signing the index of down-regulated genes as negative, followed by rank-transforming all retained genes, and dividing the genes into equal bins. Third, we used the iPAGE algorithm that calculated the mutual information between the gene ranks and the pathway memberships (the number of genes belonging to a pathway in each bin) for every Gene Ontology term. A random-permutation test was used to estimate the significance of these mutual information (MI) values so that significantly informative pathways were identified with high MI values and low p values. Finally, the hypergeometric test was used to determine whether a specific pathway was over- or under-represented in each bin. For visualization, heatmaps of pathways by bins were drawn using log-transformed p values.

### Calculation of developmental reversal index of tumor samples

To assess the level of developmental reversal for tumor samples of TCGA LIHC, KIRC, and KIRP cohorts, we asked how greatly the shift of a tumor transcriptome from a mega NAT reference (averaging gene expressions over all NAT samples of a certain cancer type) had reversed the shift of the transcriptome along the developmental trajectory of a corresponding tissue. Formally, we defined, for a sample *S*_*i*_, the developmental reversal index as:

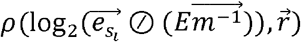

where ⊘ is element-wise division, *ρ* is the Spearman correlation coefficient, 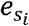 is a vector of *n* gene expressions for sample *s*_*i*_, *E* is a matrix of genes g_1_, g_2_, g_*n*_ by NAT samples s_1_, s_2_, s_*m*_ of a certain cancer type with entries as expression level, 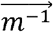 is a normalization vector of constant *m*^−1^, and 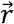 is a vector of signed goodness-of-fit values of genes g_1_, g_2_, g_*n*_ derived from the developmental RNA-seq data of a matched tissue type. We examined the association of this index with patients’ overall survival times in TCGA LIHC, KIRC, and KIRP cohorts using log-rank tests, where patients were split into two equal groups based on the median value of developmental reversal index.

### Evaluation of the utility of ECPA_cell_ as a diagnostic biomarker

To quantify the performance of ECPA_cell_ in differentiating tumors from related normal samples, we used the AUROC metric to compare it with those of all detectable individual genes (TPM ≥ 1 in ≥50% of samples in the cohort). To determine the optimal threshold of ECPA_cell_ or gene expression level for tumor-normal separation, we chose the value that maximizes Youden’s J statistic, which equals to (sensitivity + specificity − 1). If multiple optimal cutoffs existed for a biomarker whose average level was higher in NAT than in tumors, the one with the highest value was picked and vice versa.

## CRediT author statement

**Yikai Luo**: Conceptualization, Formal Analysis, Visualization, Writing – Original Draft.

**Han Liang**: Conceptualization, Supervision, Writing - Review & Editing, Funding acquisition. All authors read and approved the final manuscript.

## Acknowledgements

We thank H. Zhang, H. Chen, and other members of the Liang lab for helpful discussions. We thank H. Goodarzi for critical review of the manuscript. We also thank K. Mojumdar for editorial assistance. This work was supported by the US National Institutes of Health (U24CA209851 and Cancer Center Support Grant P30CA016672 to H.L.), an MD Anderson Faculty Scholar Award (to H.L.), and the Lorraine Dell Program in Bioinformatics for Personalization of Cancer Medicine (to H.L.).

## Authors’ ORCID IDs

0000-0001-7589-7981 (Yikai Luo)

0000-0001-7633-286X (Han Liang)

## Supplementary materials

**Figure S1.**
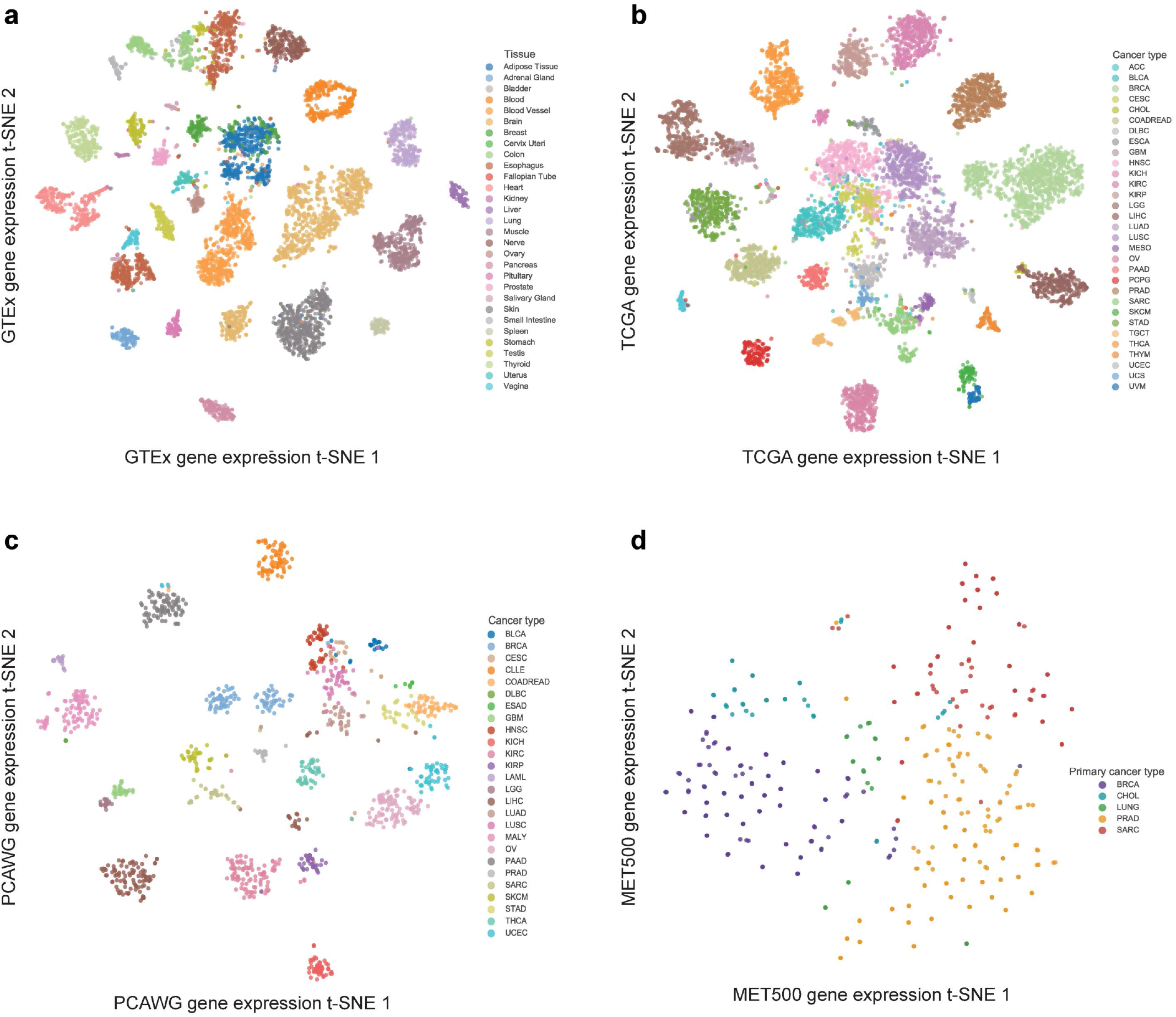
t-SNE projection of samples based on gene expression. t-SNE projection of the GTEx (**A**), TCGA (**B**), PCAWG (**C**), and MET500 (**D**) samples based on their gene expression profiles.

**Figure S2.**
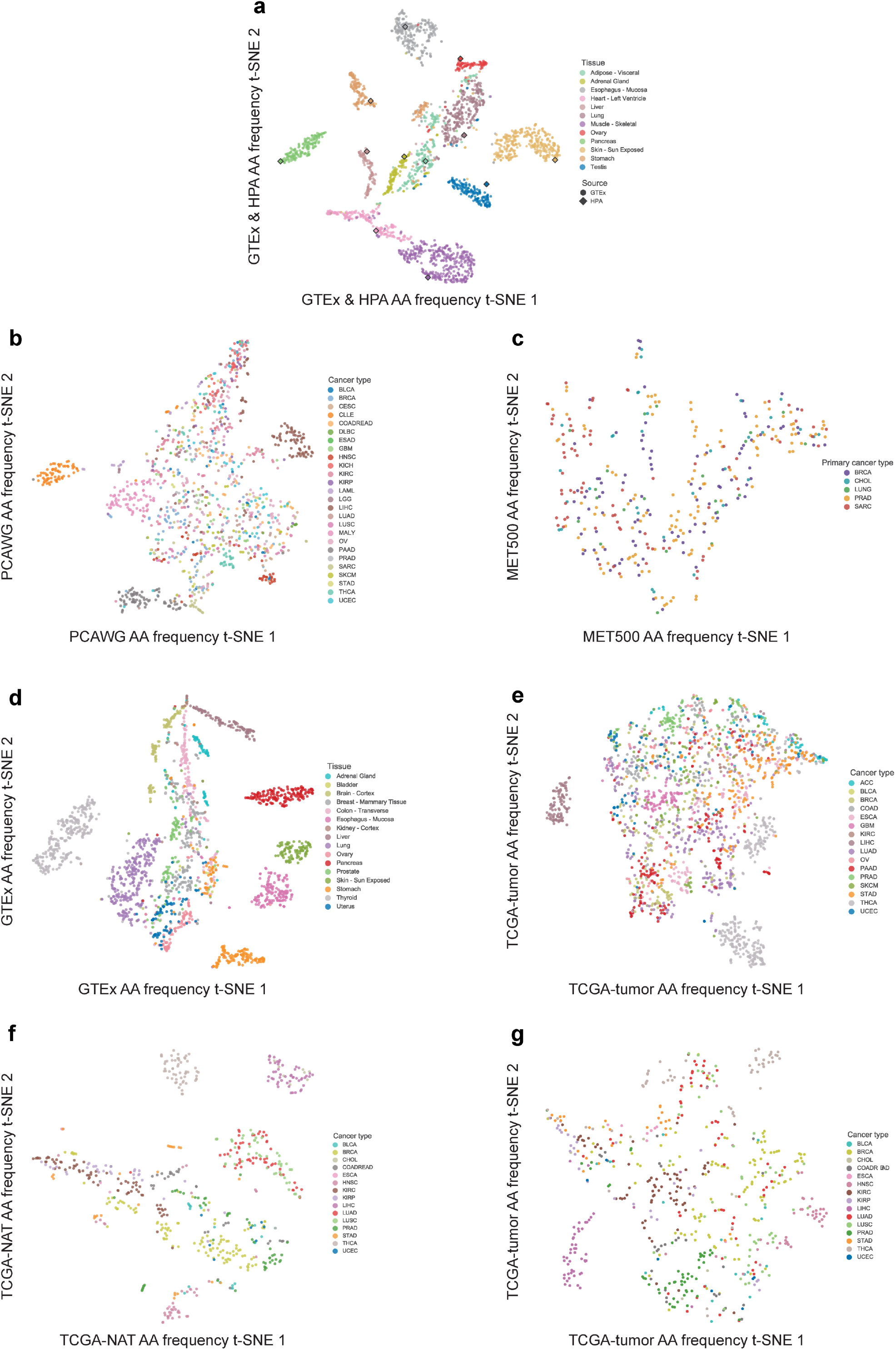
t-SNE projection of samples based on amino acid frequency. t-SNE projection of the GTEx & HPA (**A**), PCAWG (**B**), MET500 (**C**), down-sampled GTEx (**D**), down-sampled TCGA tumor (**E**), matched TCGA NAT (**F**), and matched TCGA tumor (**G**) samples based on their amino acid frequency.

**Figure S3.**
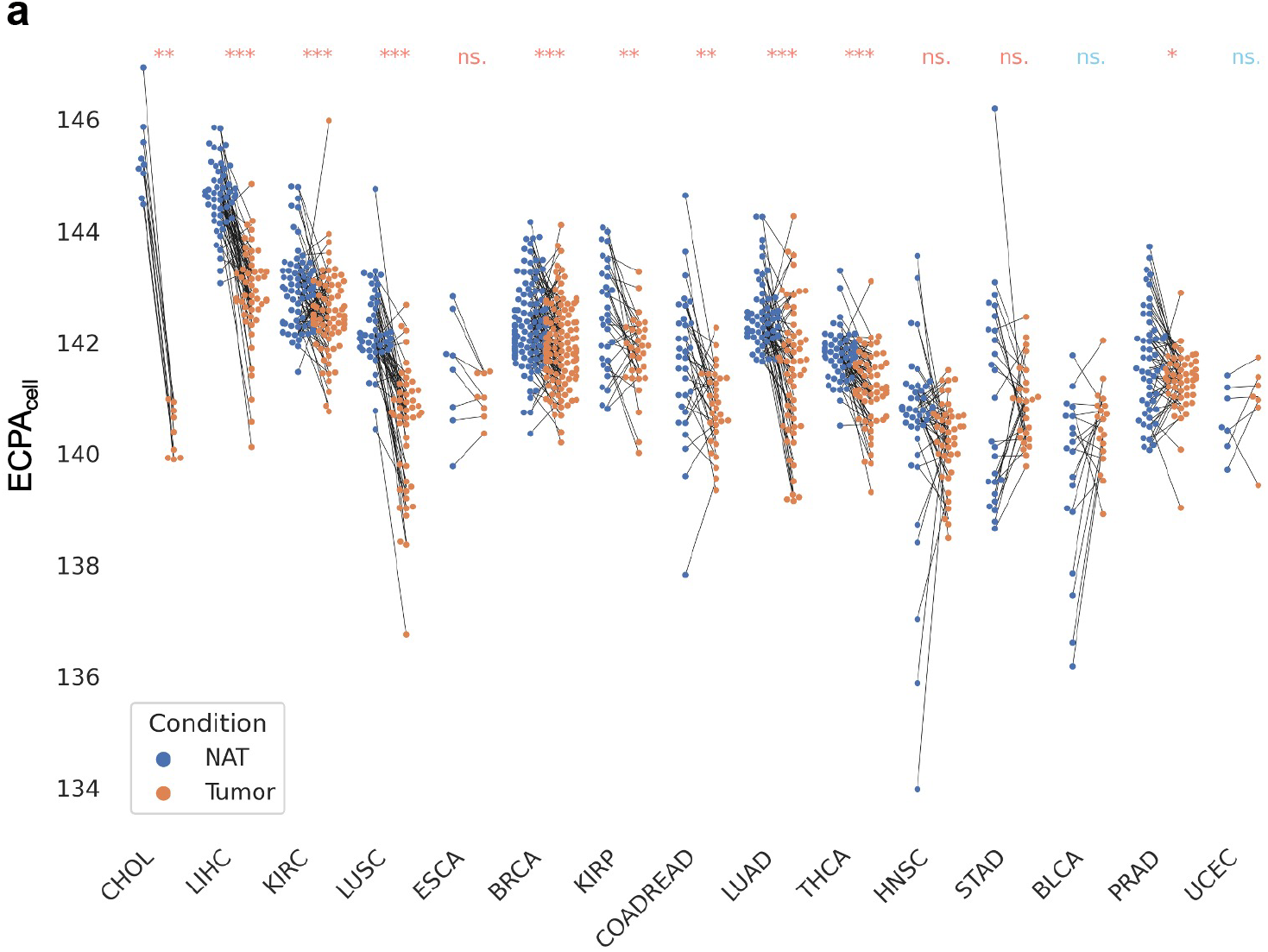
Re-calculation of ECPA_cell_ in TCGA samples without highly-expressed genes. **A**. ECPA_cell_ of tumor samples and matched normal tissue samples across TCGA cancer types after removal of genes encoding high-abundance housekeeping or tissue-specific proteins

**Figure S4.**
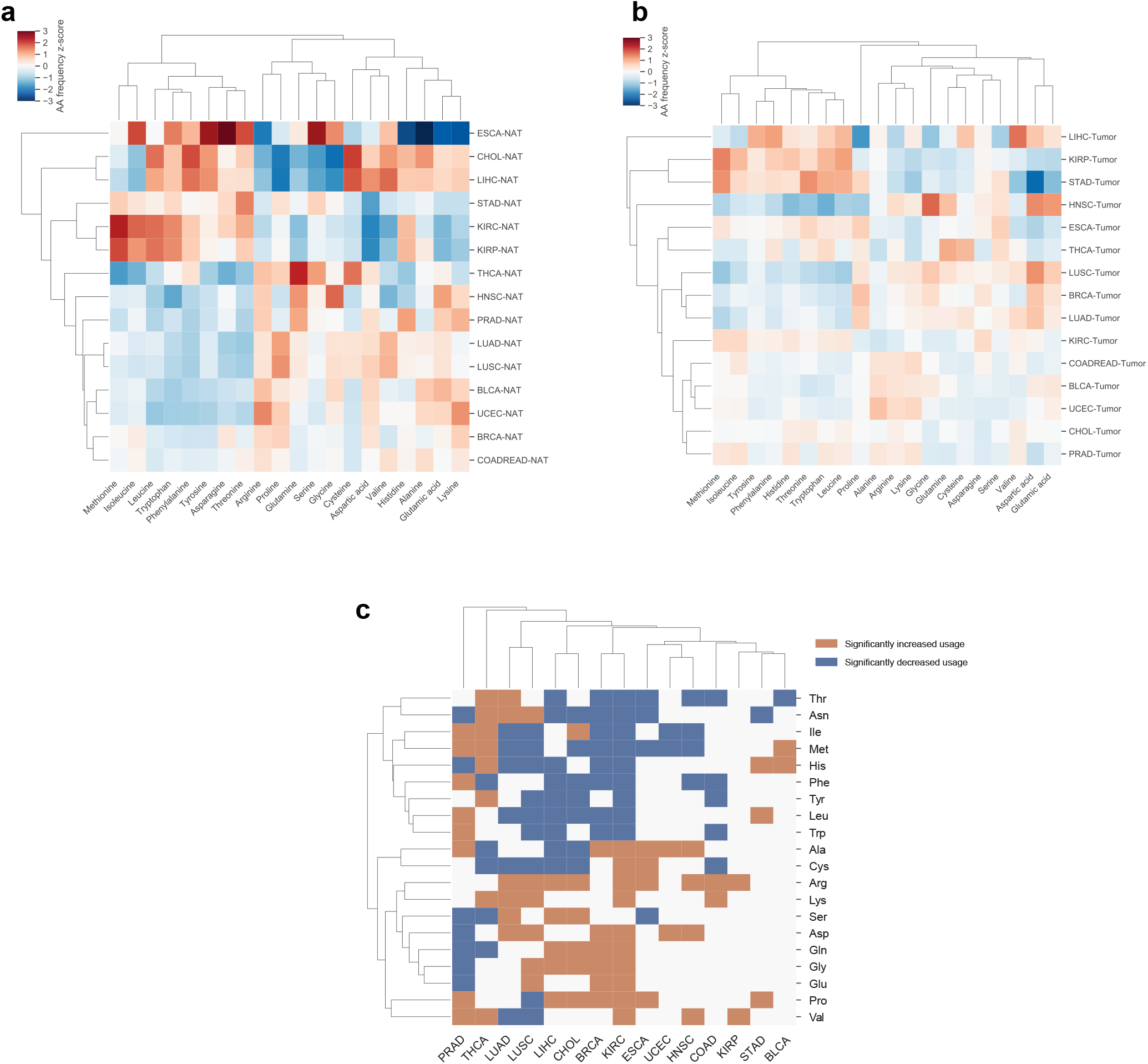
Differential amino acid usage within and between tumor and NAT samples. Heatmaps showing the average frequency of individual amino acids for NAT samples (**A**) and tumor samples (**B**), normalized as z-scores, across 15 cancer types. **C**. Heatmap showing significantly increased or decreased usage of individual amino acids between tumor and NAT samples across 15 cancer types.

**Figure S5.**
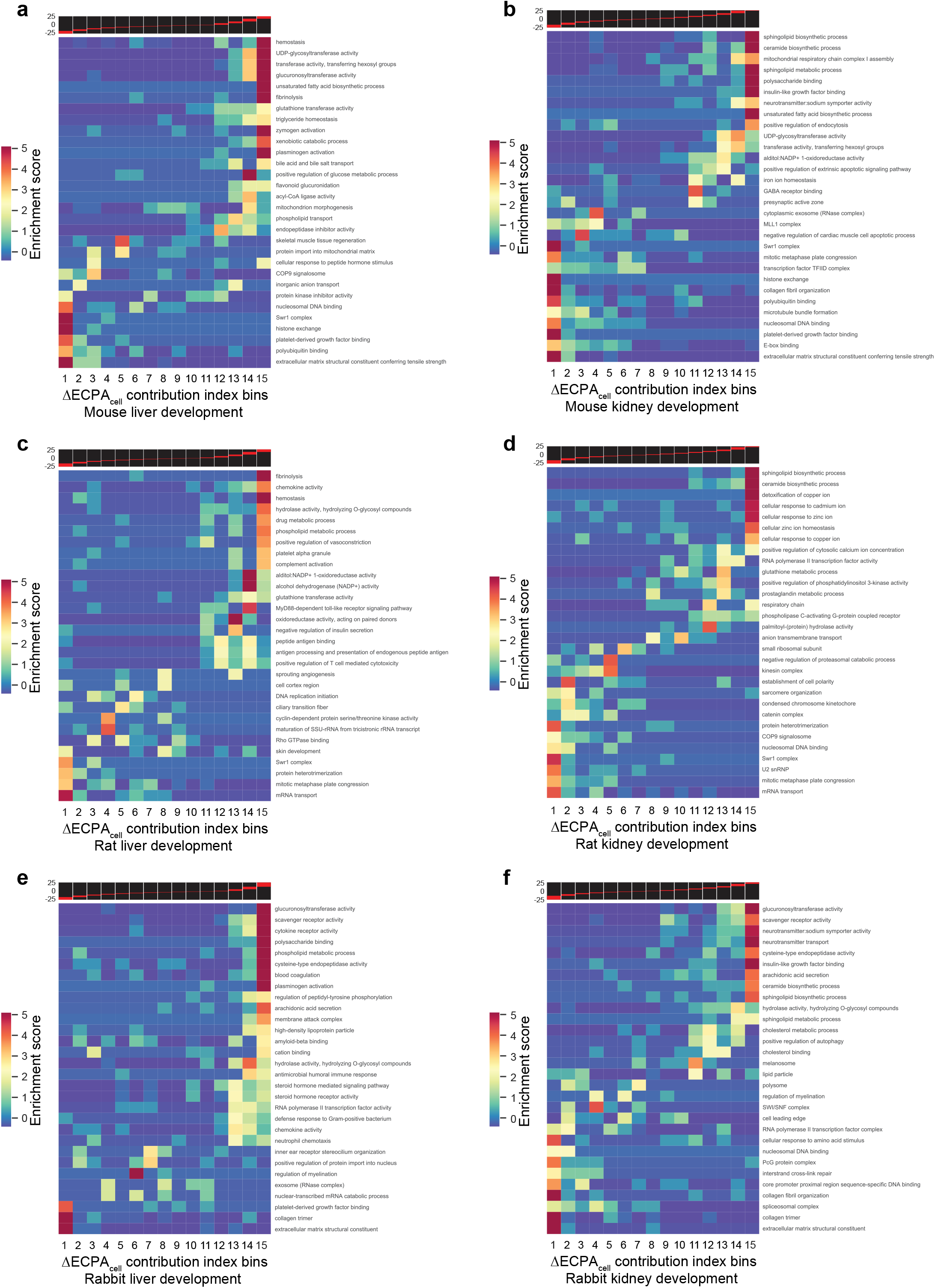
Functional enrichment of genes with a positive contribution to ECPA_cell_ in non-human tissue development. Heatmaps showing enrichment patterns of well-defined gene modules that contribute to ECPA_cell_ increase during the development of the mouse liver (**A**) and kidney (**B**), the rat liver (**C**) and kidney (**D**), and the rabbit liver (**E**) and kidney (**F**). Red stripes embedded in the black background on top of each heatmap designate the range of ΔECPA_cell_ contribution index within every bin.

**Figure S6.**
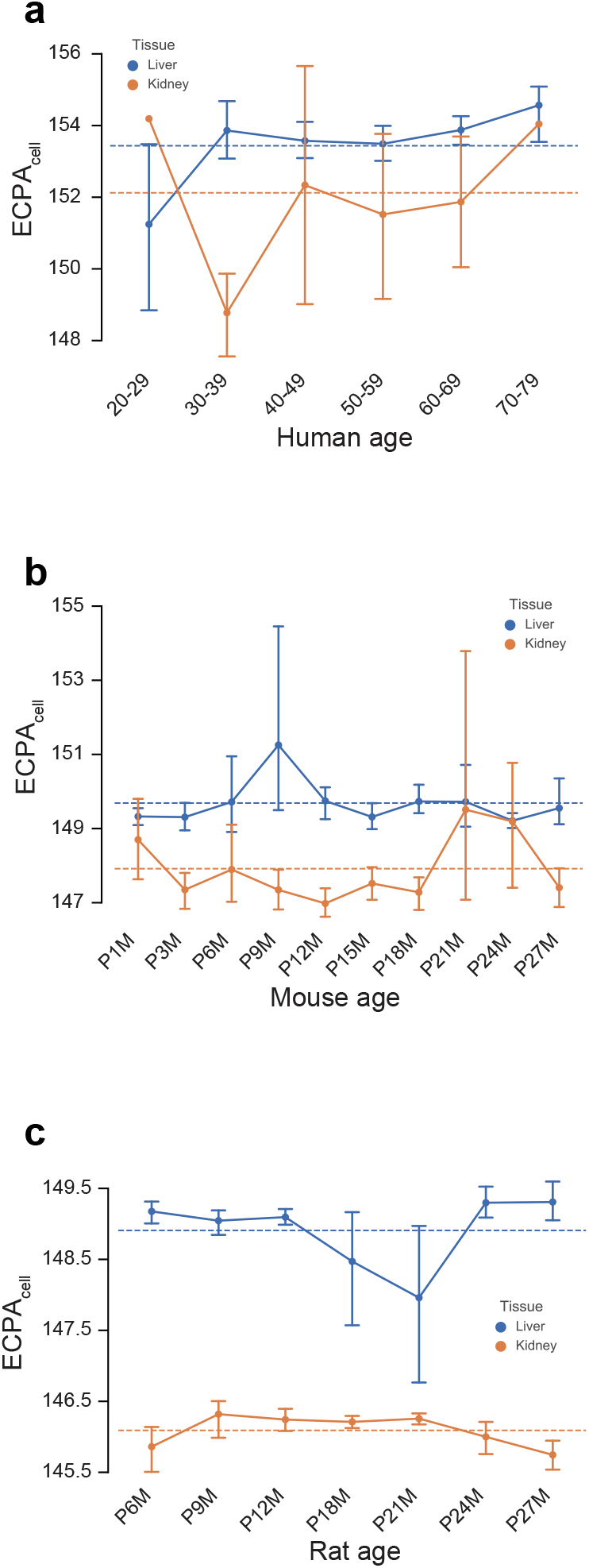
Variations of ECPA_cell_ during tissue aging. Trend lines of ECPA_cell_ during aging of the liver and the kidney in humans (**A**), mice (**B**), and rats (**C**). Blue and orange dashed lines indicate the average levels of ECPA_cell_ across age groups for the liver and the kidney, respectively. Error bars denote 95% confidence intervals.

**Figure S7.**
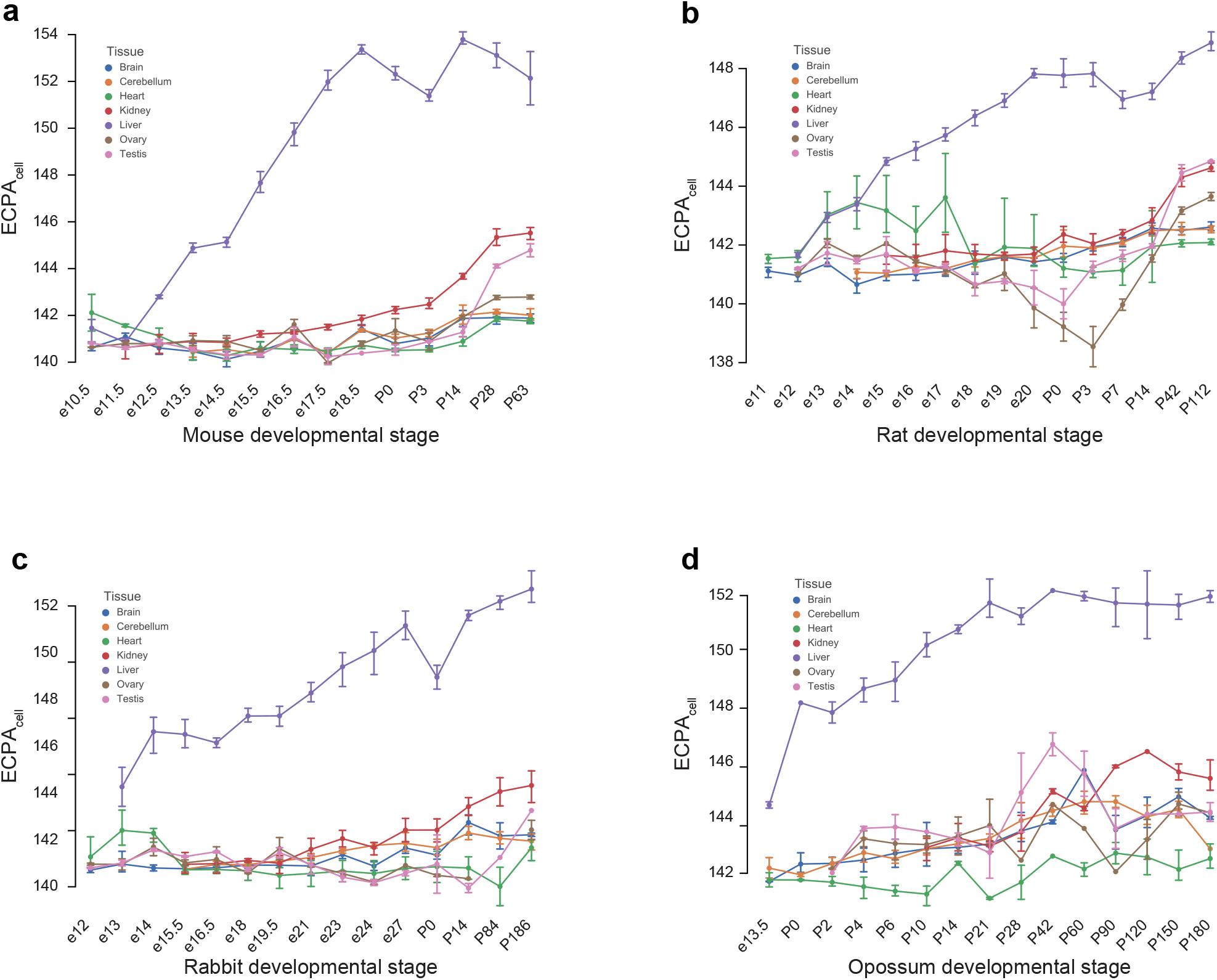
Variations of ECPA_cell_ during the development of multiple tissues in non-human mammals. Trend lines of ECPA_cell_ in seven tissues across four mammals, including mice (**A**), rats (**B**), rabbits (**C**), and opossums (**D**). Error bars denote 95% confidence intervals.

